# The fungal endophyte *Fusarium solani* provokes differential effects on the energy balance of two *Lotus* species

**DOI:** 10.1101/588400

**Authors:** Amira S. Nieva, Juan M. Vilas, Andrés Gárriz, Santiago J. Maiale, Ana B. Menéndez, Alexander Erban, Joachim Kopka, Oscar A. Ruiz

## Abstract

**Background and aims:** The interactions established between plants and endophytic fungi span a *continuum* from beneficial to pathogenic associations. The aim of this work was to explore the mechanisms underlying the potentially beneficial effects provoked by a fungal strain in legume species of the genus *Lotus*.

**Methods:** The ability to solubilise phosphorous was evaluated in nine fungal strains isolated from roots of *L. tenuis*. A selected strain was further assessed for its ability to colonize plant roots in different *Lotus* species. The effects of the two interactions were assessed by analysis of the photosynthesis, sugar amount, and macronutrient status of leaves and roots.

**Results:** A fungal isolate identified as *Fusarium solani* shows the highest phosphate-solubilisation activity and grows endophytically in roots of *L. japonicus* and *L. tenuis*. Fungal invasion enhances plant growth in *L. japonicus* but provokes a contrasting effect in *L. tenuis*. Photosynthesis, sugars and K content showed a differential effect in both plant species.

**Conclusions:** Our results indicate neither of the plant species evaluated in this work were significantly stressed by *F. solani*. Thus, the differential responses observed are due to distinct mechanisms involving photosynthesis, potassium homeostasis, and carbohydrate metabolism that are employed by plants to maintain fitness during the endophytic interaction.

## Introduction

The interactions established between microorganisms and plants have been evaluated intensively. Among them, one of the first being recognized was the mutualistic interaction established between roots and mycorrhizal fungi, a process that conduces to important improvements in plant nutrition and tolerance to stress. Subsequently, many other bacterial and fungal endophytes were reported to provide the host with several benefits (Rodriguez et al. 2009). On the other hand, the interactions between plants and endophytes may also result in pathogenesis where the fungal endophyte proliferates and causes detriment effects in the plant host (Saikkonen et al. 1998, Douglas 2010, Kiers et al. 2010). Up to date, much effort has been made to understand the fundamental mechanisms explaining the *continuum* mutualism-pathogenesis in plant-endophytic bacteria interactions, whereas those involved in the interactions established by endophytic fungi are far less known.

Much of what we know about genetics and physiology in legumes comes from studies in the model species *L. japonicus* (Handberg and Stougaard 1992). Moreover, other members of the *Lotus* genus are economically important species used in pasture systems in a diverse range of landscapes (Escaray et al. 2012). For instance, *L. tenuis* (Waldst. & Kit., syn. *Lotus glaber*; Kirkbride, 2006) is an important naturalized legume in lowland grasslands in the Flooding Pampa region in Argentina (García et al. 2008; Antonelli et al. 2016, 2019), the most important area devoted to cattle production in this country. This area is characterized by flooding and dry periods, which conduce to restrictive soil conditions such as high salinity and alkalinity as well as low fertility. Pasture promotion is a widely-used agricultural practice in the Flooding Pampa to improve the implantation of *L. tenuis*, consisting in the use of herbicides to remove weeds and promote the better growth of this species. As a consequence, the extensive cultivation of *L. tenuis* causes modification in the diversity of soil fungal communities, with an increase in the relative abundance of *Fusarium* species (Nieva et al. 2018). Therefore, it is possible to speculate that interactions between the roots of *L. tenuis* and *Fusarium* may be favoured under these restrictive soil conditions. The genus *Fusarium* is distributed in a wide range of geographical and climatic conditions, and its members establish interactions with a large group of plant hosts (Backhouse et al. 2001). Fungal species belonging to the *Fusarium* genus are the main pathogens of important crops, such as *F. graminearum* in cereals (McMullen et al. 1997), while other species, like *F. oxysporum*, are able to infect a wide variety of plant species and produce vascular wilt diseases (Di Pietro et al. 2001). Despite of the importance of *Fusarium spp*. due to the economical loses these fungi cause in important plant species, their capacity to produce symptomless infections has also been reported (Kuldau and Yates 2000). The molecular events triggered in the endophytic infections are analogous to those produced in the pathogenic ones. For example, endophytes perform microbial infections that can be detected by plants and regulated by oxidative balance and salicylic or jasmonic acid signalling pathways (Saikkonen 2013). The genus *Fusarium* has been proposed as model of soil-borne pathogens (Roncero 2003) and the mechanisms involved in their virulence has been studied (Berrocal-Lobo and Molina 2008). Nevertheless, the mechanism involved in the interactions between plant and *Fusarium* endophytes is still neglected in the literature.

The aim of this work was to isolate and select a fungal endophyte from the root system of *L. tenuis* plants growing in the Flooding Pampa soils, with ability to solubilise phosphate and to convey tolerance to alkalinity and salinity. Moreover, we characterized the effect of fungal infection on plant fitness using a selected strain and the species *L. japonicus* or *L. tenuis*. We found that phosphorous solubilising strains are also able to invade inner tissues of *Lotus* roots and to provoke differential responses in the studied plant species. These differential responses seem to be due to the deployment of distinct nutrient metabolizing mechanisms. Our results shed more light on the current knowledge of the molecular events explaining the mutualism-pathogenesis *continuum* in plant-microbe interactions.

## Materials and Methods

In order to isolate fungal endophytic strains, *L. tenuis* plants growing on Flooding Pampa soils were collected. The root system was separated from the shoots and washed under tap water. The roots were cut in several pieces of 1 cm length and placed with ethanol (70%) during 1 minute, followed by NaClO solution (10%) during 3 minutes and rinsed several times with sterile distilled water. Each root fraction was placed in potato dextrose agar medium (PDA, Britania Lab, Argentina) containing 1 mg/L gentamicin, until the apparition of fungal mycelia. The water corresponding to the last wash after the disinfection process was used as control of the surface sterilizing procedure. The strains were placed in PDA medium and storage at 4°C.

For the evaluation of the phosphate solubilisation capacity, each strain was cultivated in liquid NBRIP medium (Nautiyal, 1999) supplemented with 10 g additional of glucose, in shaker during 7 days at 30°C. After the incubation period an aliquot of the supernatant medium was collected to evaluate the amount of H_3_PO_4_ according to the Murphy and Riley (1962) procedures.

Nine strains, according to the highest phosphate solubilisation capacity were selected for further analysis. Then, they were identified through amplification and sequencing of the Internal Transcribed Spacer ITS1-ITS2 of the ribosomal 18S gene (ITS, White et al. 1990) and the alpha Transcription Elongation Factor (α-TEF, Geiser et al. 2004) gene. In order to amplify these sequences, the mix reaction (25 μl of final volume) consisted of: 1 μl of template, 1.5 μl MgCl2 (25 mM), 2.5 μl buffer reaction (10x), 0.2 μl Taq polymerase, 0.2 μl dNTP (10 mM), 0.5 μl of the primers ITS5 (5’GGAAGTAAAAGTCGTAACAAGG’3) and ITS4 (5’TCCTCCGCTTATTGATATGC ′3) (White et al. 1990), EF1 (5’ATGGGTAAGGA(A/G)GACAAGAC’3) and EF22 (5’AGGAACCCTTACCGAGCTC’3) (Geiser et al. 2004), for the amplification of ITS and α-TEF, respectively. Thermal cycles for the amplification of ITS were performed as follow: 95°C during 5 min, 38 cycles of: 94°C 30s, 53°C 30s, 72°C 45 s and a final extension of 72°C 10 min. The thermal cycles of amplification for α-TEF were performed as the following: 94°C 5 min, 38 cycles of 94°C 45 s, 56°C 30 s, 72°C 2 min, and a final extension of 72°C 2 min. In order to assign the taxonomic identity, the databases *UNITE* (version 2013, Nilsson et al. 2018) and *FUSARIUM ID* (Geiser at al., 2004) were used.

The capacity of the selected strains to growth in different concentrations of NaCl was carried out in PDA medium supplemented with 0, 100, 150 and 200 mM NaCl. The ability to growth in a wide range of pH was also determined using malt extract agar medium supplemented with buffers solutions according to the procedure described by Nagai et al. (1995). The strains were cultivated in each medium at 30°C during 7 days. At the end of the incubation period, the fungal growth was measured through the calculation of mycelial area using the *ImageJ* software v 1.47. In order to prepare fungal inoculum, the selected strain was cultivated on PDA medium at 30°C during 14 days. Then, plugs of medium were cut from the borders of the colony and used as inoculum. Plugs of fresh PDA medium were used in control plants.

### Plant growth and culture conditions

Seeds of *L. tenuis* (cv. *Nahuel*) and *L. japonicus* (ecotype *Gifu B-129*) were scarified with concentrated H_2_SO_4_ and washed ten times with distilled water. Then, seed surface was sterilized with NaClO (5%) and rinsed several times with sterile distilled water. Sterilized seeds were incubated in sterile water overnight and placed in Petri dishes containing water-agar medium (0.5%). Plates were incubated during 15 days until the radicle and trifoliate leaf developed. The germination was carried out in a grow chamber, with a 16/8 h photoperiod at 24°C/19°C (day/night), light intensity of 240 mol m^-2^s^-1^ and 60% humidity. Ten seedlings were grown on pots containing sterile sand-perlite substrate (2:1) and irrigated with nutritive *Evans* solution (Evans 1970). The inoculation with the selected strain was carried out immediately after transplant. Seedlings were cultivated during 37 days as described above. After this period, roots were washed under tap water and separated from the shoots. An aliquot of the roots were surfaced sterilized in order to re-isolate associated fungal strain as a way to check its endophytic nature, using the same procedures explained above. Fractions of roots were stained with Trypan blue according to the procedures explained by Phillips and Hayman (1970) for microscopic analysis. Each plant fraction (roots and shoots) was placed at 70°C during 48 h for dry weight assessment. Foliar area was measured in the second oldest leaf of main shoots.

### Net photosynthesis rate and stomatal conductance

Carbon exchange was determined through an infrared gas analyser (Portable Photosynthesis System, MA, USA) supported with LED lights with an intensity of 240 mol m^-2^s^-1^, according to the procedures described by Calzadilla et al. (2016). Net photosynthesis (PN), CO_2_ in the sub-stomatal cavity (Ci) and stomatal conductance (GS) were calculated through the analysis of the parameters obtained from the infrared gas analyser.

### *Chlorophyll a* transient fluorescence analysis

*Chlorophyll a* fluorescence transients were used to characterize the status of the photosystem II (PSII), using a Portable Plant Efficient Analyzer (PEA, Hansatech Instruments Ltd., UK). With this purpose, leaves were adapted to dark before measurement during 20 minutes and exposed to a light intensity of 3500 μmol m^-2^s^-1^ during 3 s. Parameters derivate from *OJIP* test (O: Initial fluorescence, at 50 μs or less, J-I: intermediate levels at 2 ms and 30 ms, P: maximal fluorescence) were analysed in order to detect the variations in the PSII status. In order to characterize the performance of the PSII under biotic stress, the different parameters as explained by Vilas et al. (2018) were analysed: Fv/Fm (Φ_Po_): Maximum quantum yield of the primary photochemistry and Performance Index on the Absorption Basis (PIabs), contribution of the absorption of light energy (ABS), normalized total complementary area above the *OJIP* transient (Sm), trapping of excitation energy (TRo), conversion of excitation energy to electron transport (ETo), Dissipation flux (DIo), per reaction centre (RC) and Cross section (CSo), Ψ_Eo_: Electron transport from *quinone a* to plastoquinone efficiency, Φ_Eo_: maximum yield of electron transport from *Qa* to plastoquinone primary photochemistry.

### Determination of photoassimilates

The amount of fructose, glucose, sucrose and maltose was determined through gas chromatography coupled to electron impact ionization/time-of-flight mass spectrometry (GC-EI/TOF-MS, Agilent 6890N24, Agilent Technologies, Böblingen, Germany). Samples of leaves and roots were collected from inoculated or mock-inoculated plants and immediately deep frozen. The polar metabolites were obtained through chloroform: methanol extraction, according to previously reported procedures (Lisec et. al 2006) with modifications described in detail by Dethloff and co-authors (2014). GC-EI/TOF-MS chromatograms were processed through a standardized data analysis procedure using ChromaTOF (v. 4.32; LECO, St. Joseph, USA) and TagFinder software (Luedemann et al. 2008). Compounds were identified by mass spectral and retention time index matching to the reference collection of the Golm metabolome database (GMD, http://gmd.mpimpgolm.mpg.de/; e.g. Hummel et al. 2010) and to the mass spectra of the NIST08 database (http://nistmassspeclibrary.com/).

### Macronutrients quantification

Total nitrogen (N) was quantified in 50 mg of dry material from leaves. Samples were digested in concentrated H_2_SO_4_ according to the Kjeldahl method adjusted for plant determinations (Bremner et al. 1982).

For Total phosphorus (P) and potassium (K) measurements, samples were digested with HNO_3_ (Merck, Germany) and diluted in 50 ml of distilled water. The macronutrients were quantified directly with a Microwave Plasma–Atomic Emission Spectrometry analyzer (MP-AES Agilent Technologies, Germany).

## Results

### Characterization of fungal strains isolated from the roots of *L. tenuis*

We evaluated the phosphate solubilisation capacity of 9 fungal strains isolated from healthy roots of *L. tenuis* plants growing in Flooding Pampa soils. All the strains were able to solubilise phosphate *in vitro*. Among them, the strains *142L52A* and *142L52B* showed a remarkable activity for phosphate solubilisation (Fig. 1). These fungal strains were identified as *Fusarium oxysporum* and *F. solani*, respectively, by sequencing the ITS region and TEF gene and comparing these data at the UNITE and Fusarium ID databases (Supplementary material). Moreover, as *L. tenuis* usually thrives in areas from the Flooding Pampa dominated by high salinity and alkalinity conditions, we evaluated the capacity of the two fungal strains to growth in a wide range of pH and NaCl concentrations. These experiments showed that *F. solani 142L52B* grows at normal rates in saline concentrations up to 200 mM NaCl and at pH up to 10, respectively. Therefore, we selected this strain for further experiments (Fig. 2).

**Fig. 1.**
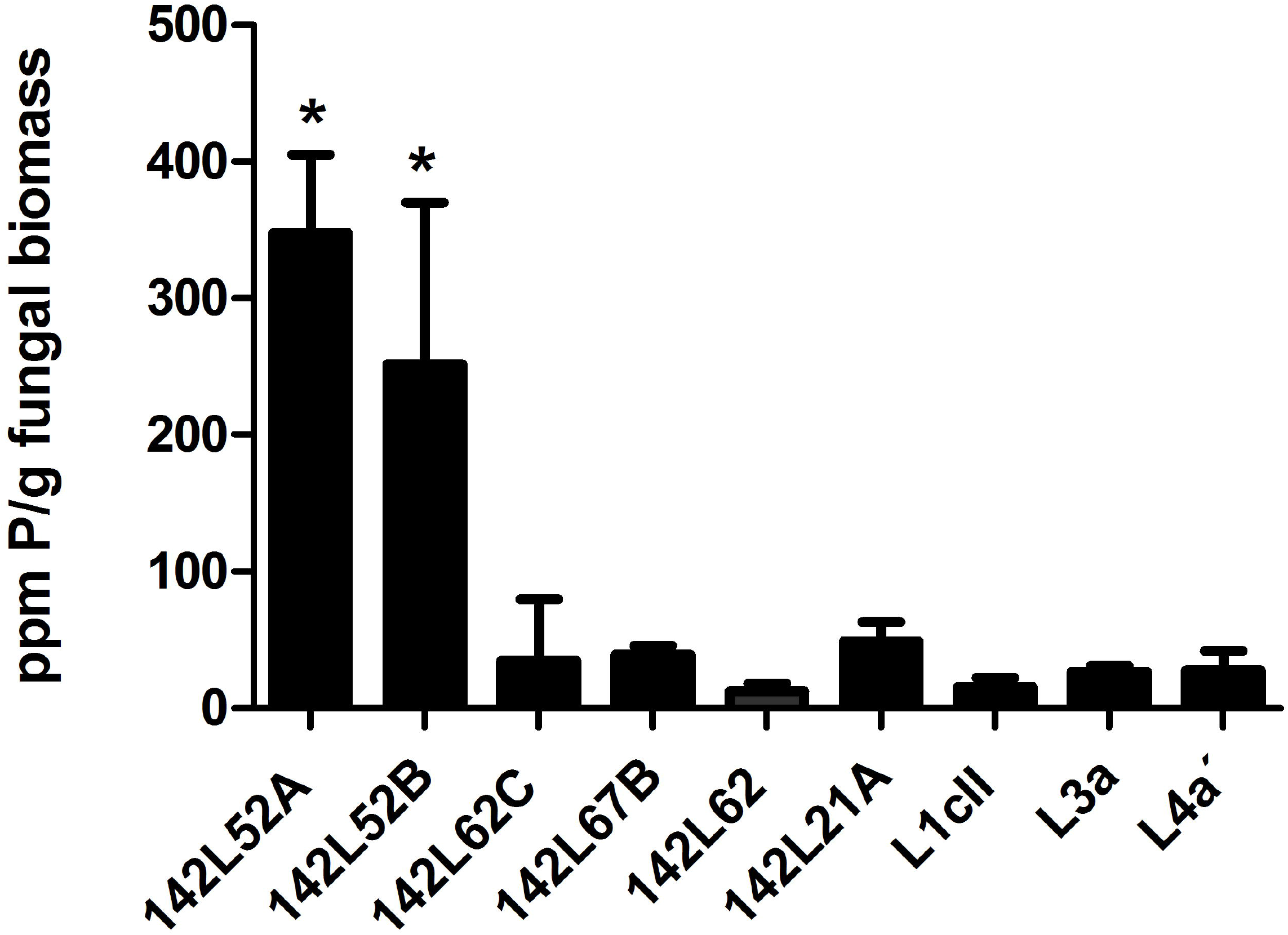
Phosphate solubilising activity of nine fungal strains isolated from healthy roots of *L. tenuis* plants growing in alkaline al saline soils. Phosphate (P) was determined after 7 days of fungal growth on NBRIP medium by blue molybdenum technique. Five biological replicates were measured (mean± standard deviation). * p<0.05 Tukey test.

**Fig. 2.**
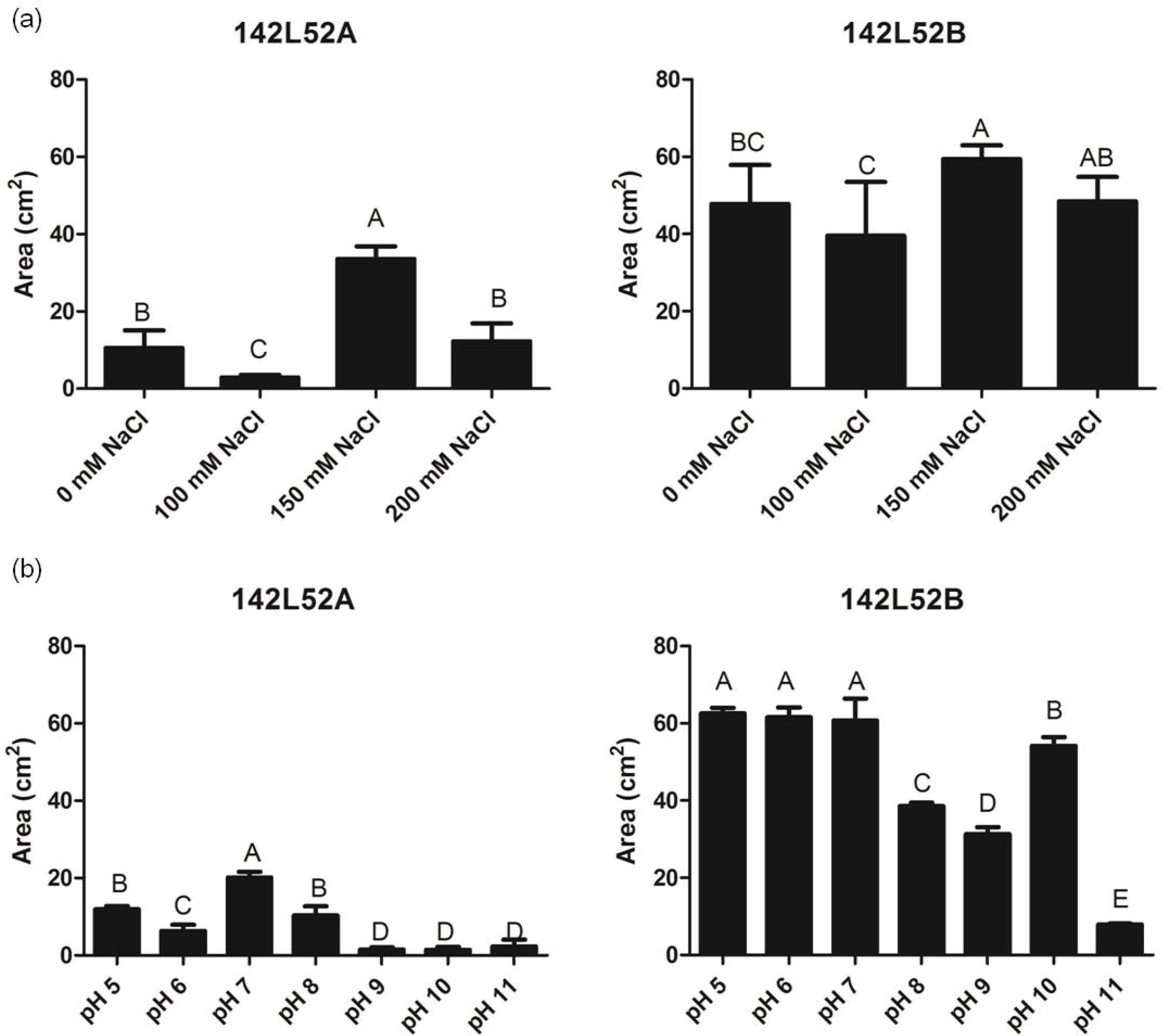
Growth of two selected strains on: a) PDA medium supplemented with either no additional NaCl (0 mM) or, 100 mM NaCl, 150 mM NaCl and 200 mM NaCl; b) Malt medium (MA) supplemented with buffer solutions, according to Nagai (1997) in order to obtain media ranging from pH 5 to 11. Fungus area was measured in five biological replicates (mean ± standard deviation), after 7 days of growth. Different letters indicate conditions that are significantly different according to the Tukey Test (p<0.05)

### Effects of *F. solani 142L52B* infection on plant growth

In order to reveal the effects of *F. solani 142L52B* on *Lotus* species, we inoculated *L. tenuis* and *L. japonicus* seedlings with mycelia plugs and incubated the plants for 37 days in a growth chamber. As shown in Fig. 3a, fungal inoculation caused contrasting effects on the growth of the *Lotus* species evaluated in this work. In this trend, *F. solani 142L52B* enhanced growth in *L. japonicus*, whereas an impairment of growth was observed in *L. tenuis*. This effect was consistent with the changes of the biomass of roots and shoots between infected and control plants (Fig. 4a, b). Interestingly, even though the total number of leaves and lateral shoots did not change in both species in response to fungal infections (Fig. S1), fungal infection provoked a reduction in the foliar area of *L. tenuis* (Fig. 4c).

**Fig. 3.**
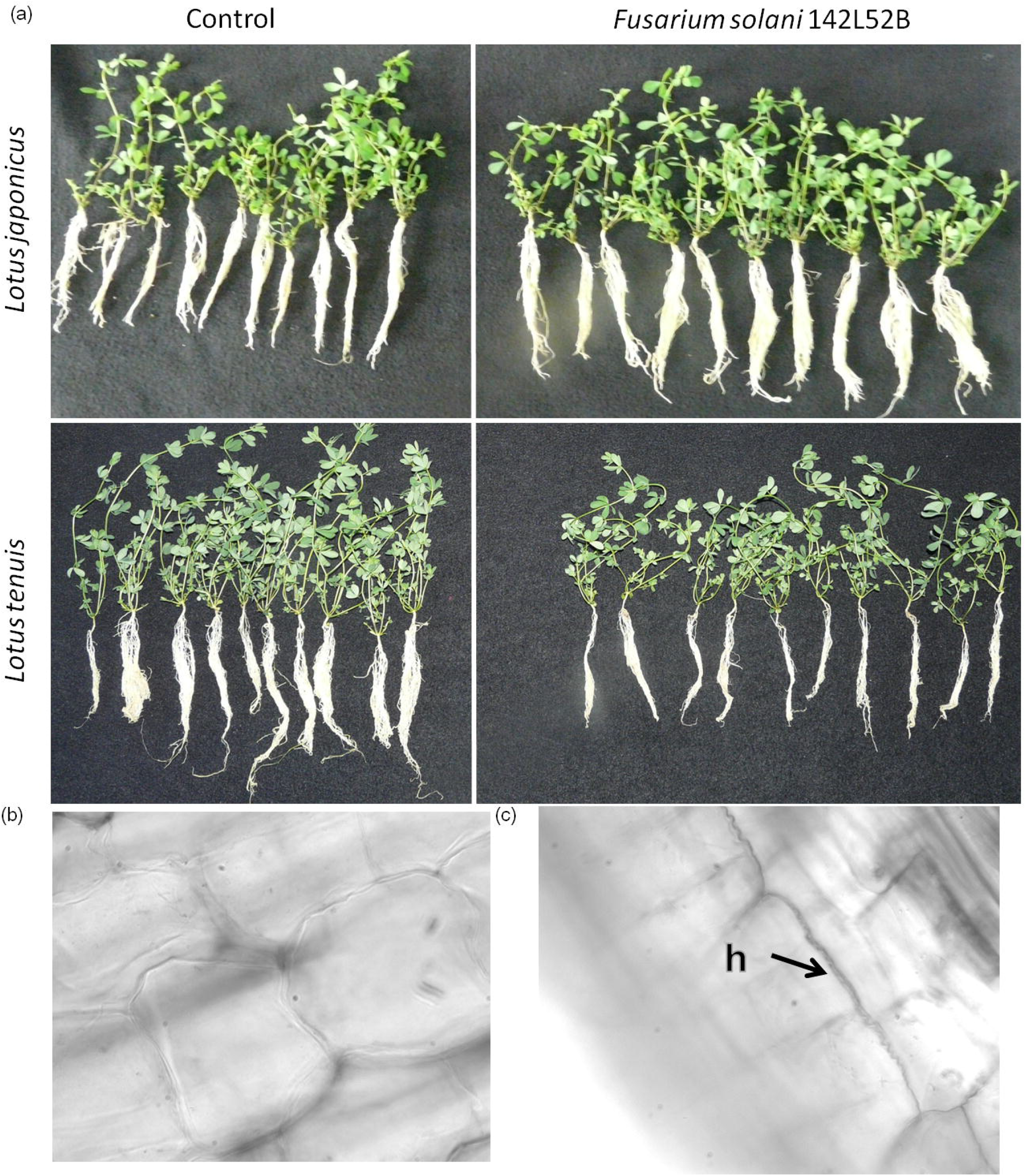
Control and *F. solani*-infected plants after 37 days of incubation. Seedlings were transferred to pots containing sterile sand-perlite (2:1). Roots were inoculated with plugs of non-inoculated PDA medium (Control) and plugs that were inoculated with *F. solani* 142L52B. (a) Whole plants after the growth period. (c). Cortex cells of roots of control plants (400x). (c) Root cortex cells after *F. solani* 142L52B infection. h: hyphae infecting the intercellular space of root cortex cells.

**Fig. 4.**
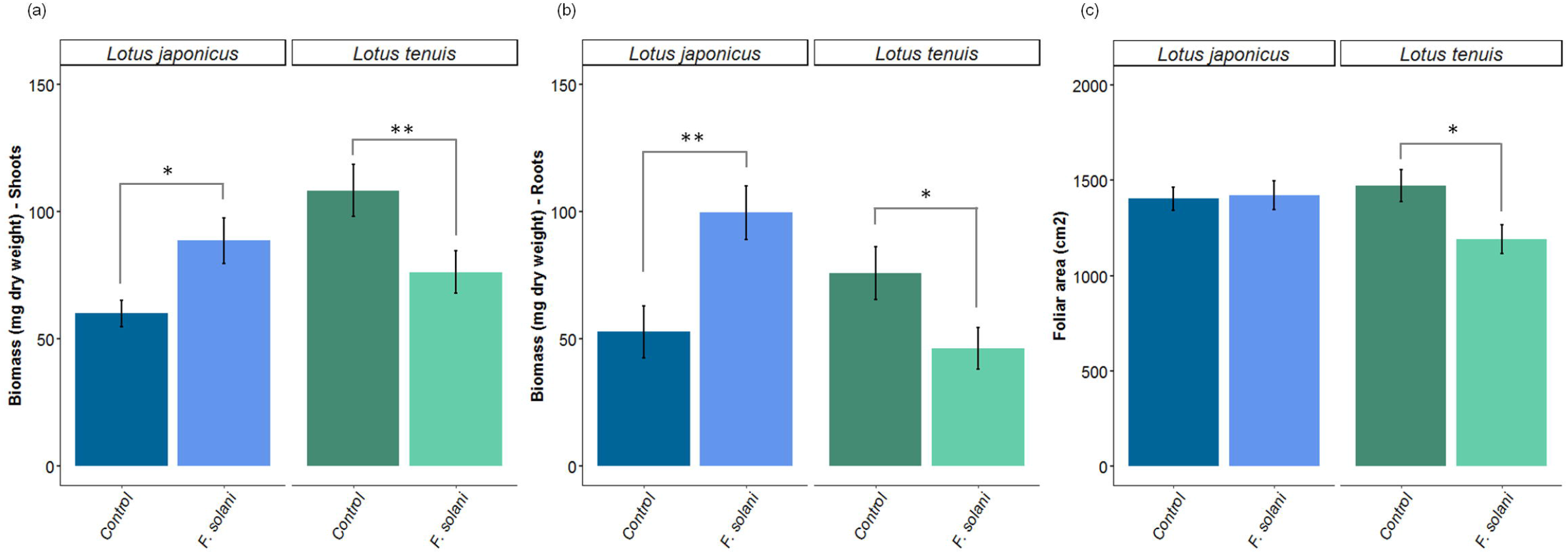
*F. solani 142L52B* provokes contrasting changes in biomass production in *L. tenuis* and *L. japonicus*. Biomass of shoots (a) and roots (b) of *L. japonicus* (blue) and *L. tenuis* (green) plants under mock and fungal-inoculated conditions. (c) Foliar area of the second oldest leaf belonging to the main shoot of *L. japonicus* and *L. tenuis* plants expressed as cm^2^. Statistical differences between controls and *F. solani* treatments were assessed using Student’s test (mean± standard deviation, n=10). * (p<0.05), ** (p<0.01).

In addition, with the purpose to confirm the endophytic nature of the fungal strain, we re-isolated it from infected roots after the growth period of 37 days. This analysis demonstrated that *F. solani 142L52B* was absent in the basal shoots and limited to the root system. Besides, microscopy analysis showed that the infection is intracellular and located only to the cortex cells of roots (Fig. 3b, c).

### Effect of *F. solani* on the photosynthesis and stomatal conductance of *Lotus* plants

The photosynthesis is sensitive to several biotic stress conditions. Because the fungus may constitute an additional carbon sink, we speculated that the endophyte could affect the photosynthetic rate. In addition, according to other studies, the endophytes could also affect stomatal opening by maintaining turgor pressure in leaves (Richardson et al. 1993). Our results showed that the net photosynthesis rate (PN), measured by CO_2_ exchange is significantly improved in *L. japonicus* infected by *F. solani 142l52B* compared to mock-inoculated controls (Fig. 5a). In turn, PN was not affected in *L. tenuis* plants by fungal colonization. The increment of PN in *L. japonicus* is in agreement with the decrease in the amount of CO_2_ in the sub-stomatal cavity (C_i_) (Fig. 5c), which reflects a higher movement of CO_2_ to the leaf mesophyll that is tightly linked to C fixation. In turn, infection by *F. solani 142L52B* did not provoke any alteration in the stomatal conductance (GS) in both plant species (Fig. 5b).

**Fig. 5.**
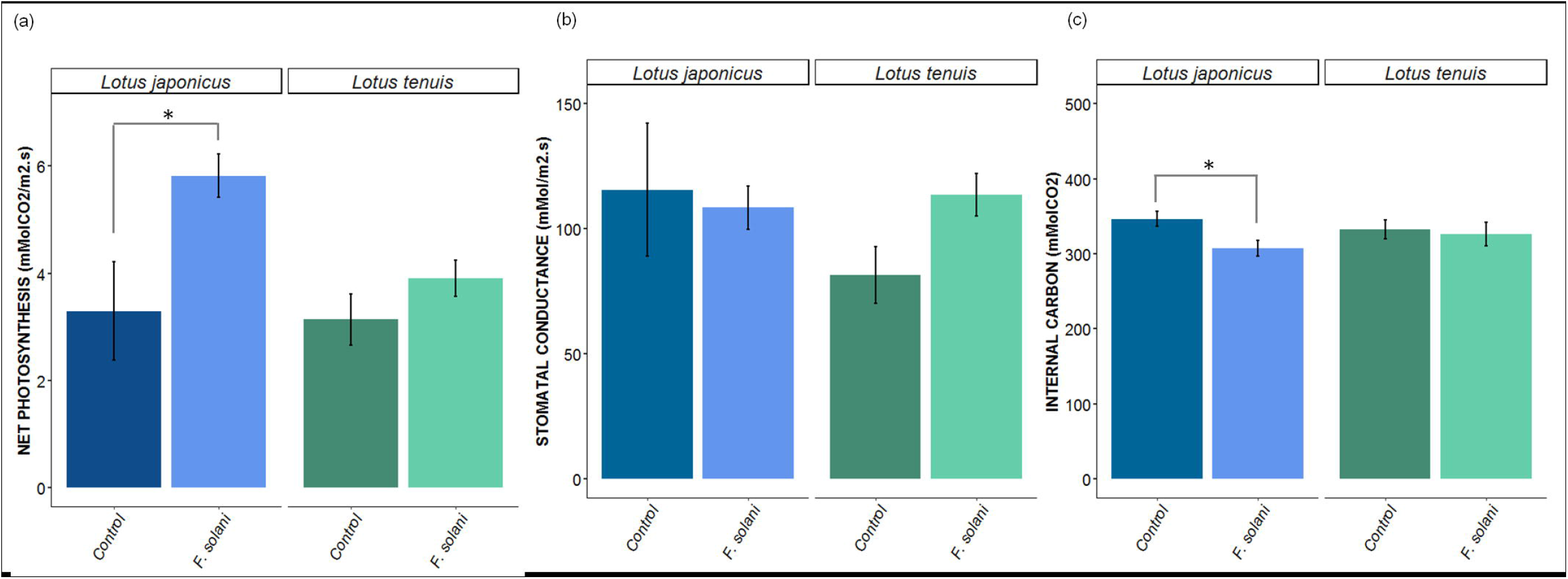
a) Net Photosynthesis (PN), b) Stomatal conductance (GS) and CO_2_ in the sub-stomatal cavity (C_i_) measured after 37 dpi (days post inoculation) in ten control treated and ten *F. solani* inoculated plants of *L. japonicus* and *L. tenuis*. Statistical differences between controls and *F. solani 142L52B* treatments were assessed using Student’s test (mean ±standard deviation, n=10). * (p<0.05), ** (p<0.01).

In addition, we analysed the Chlorophyll *a* fluorescence *OJIP* transient to determine the operational status of the photosynthetic apparatus. This non-invasive method provides valuable structural and functional parameters associated to the plant photosynthetic machinery (Berger et al. 2007). The *OJIP* test indicated that the maximum quantum yield of primary photochemistry, Fv/Fm of both plant species is not affected by the fungal infection (Fig. 6). In addition, the values of Fv/Fm for all the treatments were approximately 0.8, values that normally occur in non-stressed plants. In turn, the values of performance index (PIabs) decreased in *L. tenuis* under the fungal infection.

**Fig. 6.**
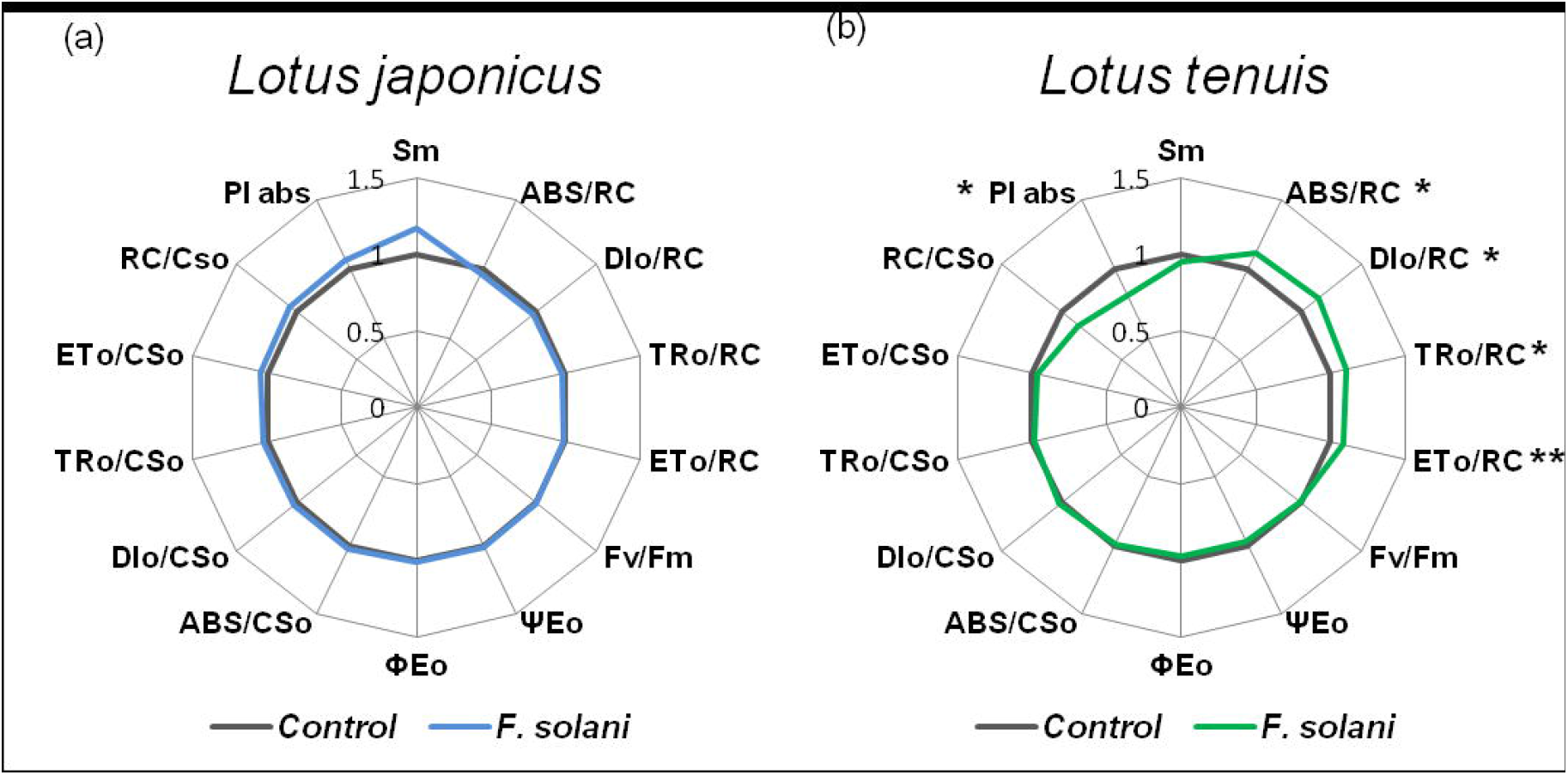
Ratios of *F. solani 142L52B /* control of each parameter obtained from an *OJIP* analysis of *L. japonicus* and *L. tenuis*. Sm: Normalized total complementary area above the *OJIP* transient PIabs: Performance index on the absorption aasis, ABS: Absorption flux, DIo: Dissipation flux, TRo: Trapped energy flux, ETo: Electron transport flux, RC: Reaction centre, CSo: Cross section, Fv/Fm (Φ_Po_): Maximum quantum yield of primary photochemistry, Ψ_Eo_: Electron transport from *quinone a* to plastoquinone efficiency, Φ_Eo_: maximum yield of electron transport from *Qa* to plastoquinone primary photochemistry, RC/CSo: active reaction centres per cross section. Statistical differences of the controls compared to *F. solani* treatments of each OJIP-derived parameter were assessed using Student’s test. * (p<0.05), ** (p<0.01).

In addition, PIabs, CSo-related and RC-related parameters were also significantly modified in *L. tenuis* under the fungal infection (Fig. 6b).

### Changes of the content of photoassimilates and ions in response to fungal infection

Metabolites such as sugars are good reporters to determine the potential cost of the interaction generated by the fungus, as their concentration is a proxy of the plant energetic balance during the endophytic interaction. As shown in figure 7, the amount of fructose, glucose, sucrose and maltose in leaves were significantly higher in *L. japonicus* infected with *F. solani 142L52B* than in controls (Fig. 7). In contrast, fungal infections led to a reduction in the concentration of these compounds in *L. tenuis*. In agreement with the increase of glucose and fructose, the amount of precursors, such as fructose-6-phosphate and glucose-6-phosphate varied in both species.

**Fig 7.**
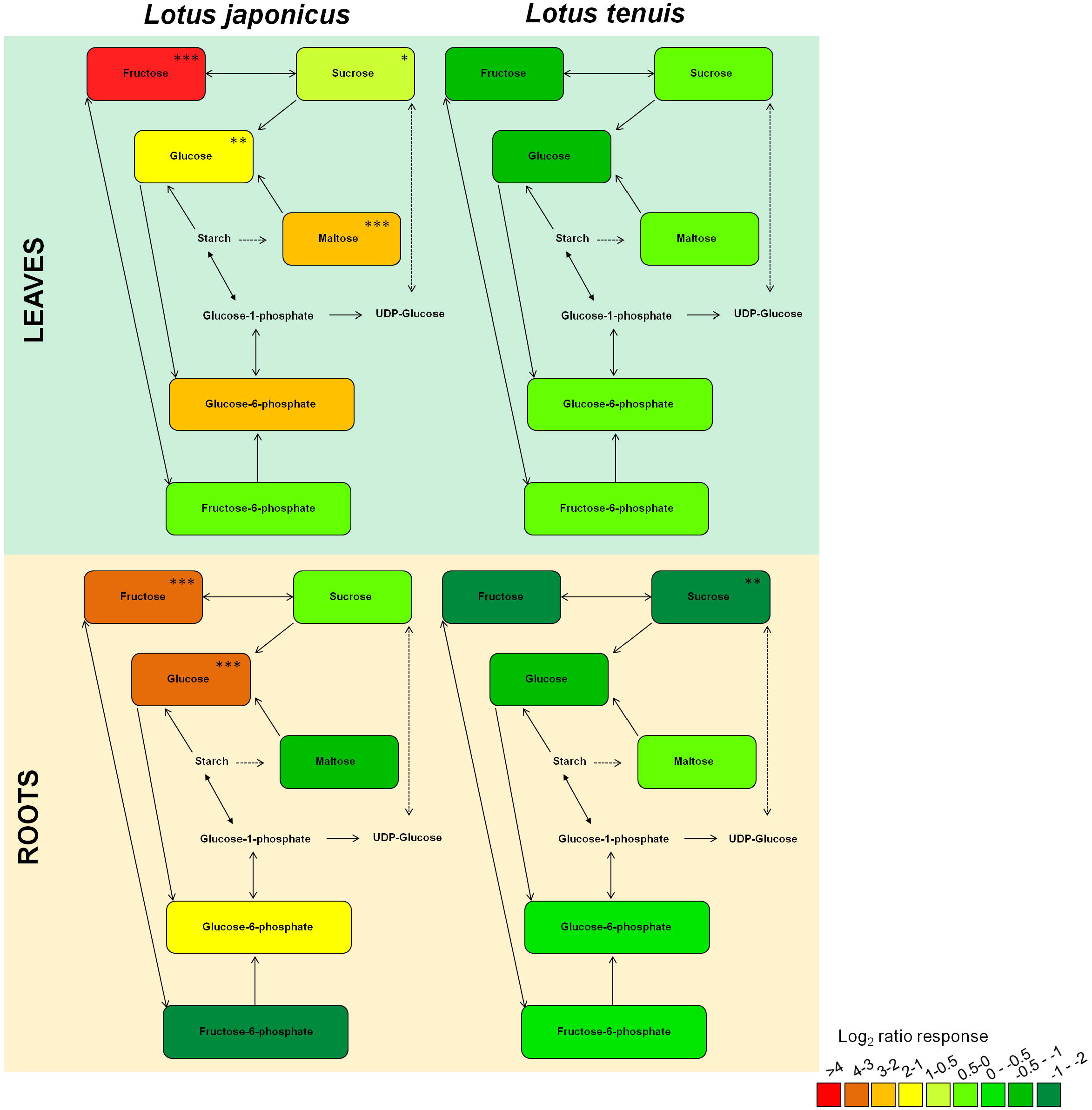
Log_2_-transformed relative sugar concentrations of *F. solani 142L52B* -inoculated versus mock-inoculated control plants. Sugars were determined by GC-EI/TOF-MS based metabolite profiling in paired leaf and root samples of *L. japonicus* and *L. tenuis*. Log2-transformed ratios are colour-coded in the range of >4 (red) to −2 (dark green) according to the inserted scale. Statistical differences between controls and *F. solani* treatments were assessed using Student’s test on ten replications per treatment. * (p<0.05), ** (p<0.01), *** (p<0.0001).

The concentrations of sugars detected in roots were also modified in both species. Thus, in *L. japonicus*there was a significant increment in the concentration of glucose and fructose, while sucrose concentration was diminished in *L. tenuis* (Fig. 7).

Fungal endophytes may provide nutritional benefits to the plant host, which could be due attributed to a better acquisition of macronutrients. Considering the ability of the strain to solubilise phosphate, we assessed the amount of P in leaves of *L. japonicus* and *L. tenuis* in order to evidence a positive effect of the endophyte in the incorporation of this element. In addition, the amount of other essential macronutrients, such as N and K were included in our analysis as it has been reported that these may be modified during the interaction with other endophytes, like *Acremonium* in tall fescue (Lyons et al. 1990) and *Peniciullium* in *Aegiceras corniculatum* (Xu et al. 2007). The effects of the fungal infection on macronutrient contents were not statistically different for N and P (Fig. 8a, b). These results indicate that there is no apparent benefit provided by the endophyte on the P and N nutrition relative to accumulated dry mass. However, a reduction in the amount of K in leaves of *L. japonicus* was observed, while an increment in the concentration of this element was detected in *L. tenuis* after the infection by *F. solani* (Fig. 8c).

**Fig. 8.**
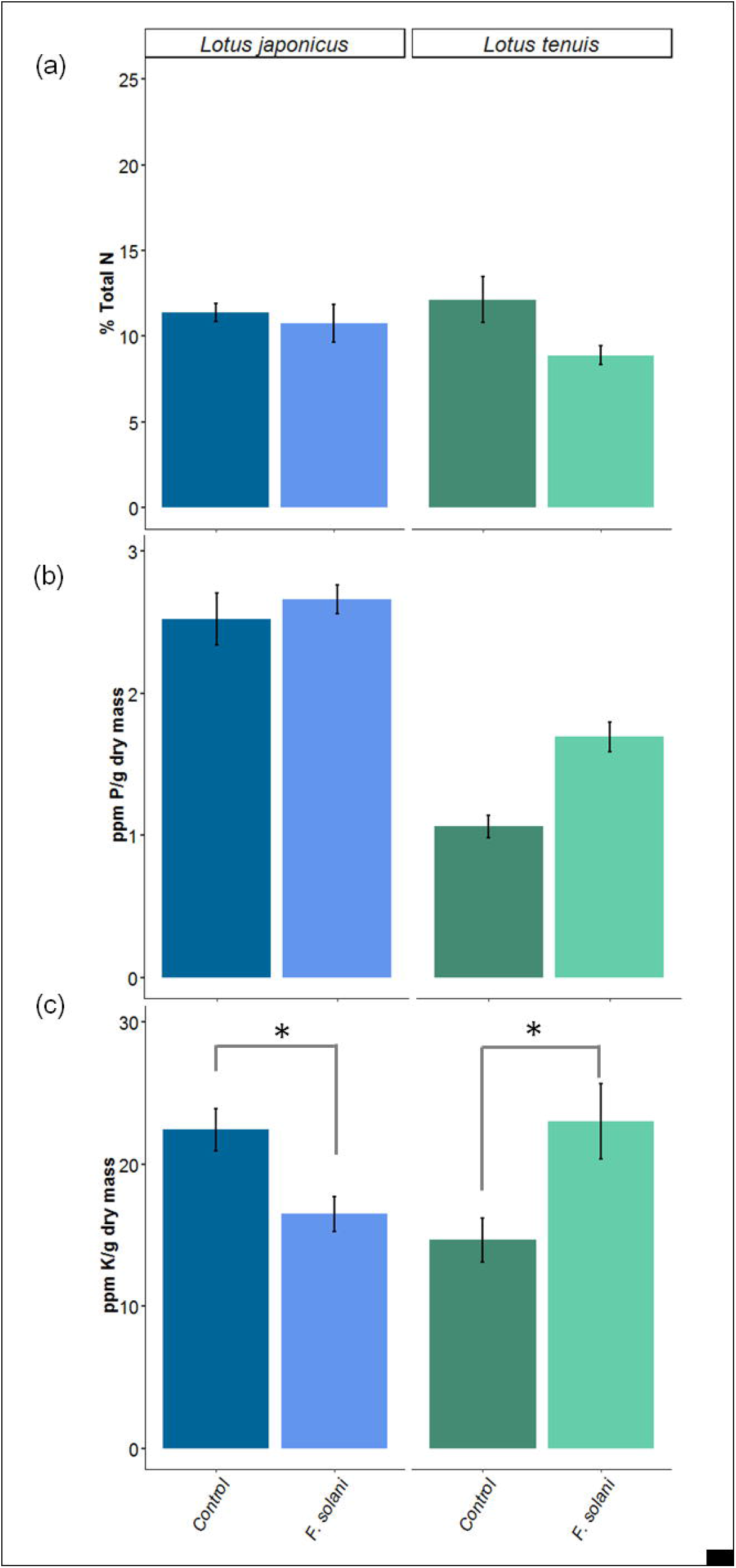
The *F. solani* effect on macronutrients measured in leaves of *L. japonicus* and *L. tenuis*. Percent of total nitrogen, N (a), concentration of phosphorous, P (b) and potassium, K (c) in leaves of *L. japonicus* and *L. tenuis* 37 days after inoculation with *F. solani 142L52B*. Statistical differences of the parameters measured were evaluated using Student’s test (mean± standard deviation, n=10). * (p<0.05).

## Discussion

In this study, we isolated and identified a strain of *F. solani* from healthy roots of *L. tenuis* growing in saline-alkaline lowland soils of the Salado River basin (Argentina), able to solubilise phosphate and tolerate alkalinity and salinity *in-vitro*. *F. solani* is an important pathogen affecting crops belonging to the *Solanaceae* family, like *Solanum tuberosum, S. Melongena, Lycopersicon esculentum* and *Capsicum anuum* (Romberg and Davis 2007). This fungal species is also able to infect other plant families, including legumes, such as *Pisum sativum* and *Glycine max*, causing the so-called sudden death syndrome (Matuo and Snyder 1973, Burke and Hall 1991, Ruan et al. 1995). Despite of this, it has been also reported that strains of this fungal species survive without causing apparent negative effects as endophyte in different plant species such as *Camptotheca acuminata* (Kusari et al. 2009), *Lycopersicon esculentum* (Kavroulakis et al. 2007, Imazaki and Kadota 2015), *Taxus baccata* (Tayung et al. 2011) and *Apodytes dimitata* (Shweta et al. 2010). To our knowledge, the present work constitutes the first report demonstrating the endophytic relationship between members of the plant genus *Lotus* and *F. solani*. These results agree with previous studies on the fungal species infecting the native flora in mediterranean areas in Europe, which indicates the presence of *F. avenaceum* as endophyte of healthy plants of *L. creticus* growing in salty soils (Maciá-Vicente et al. 2008). Thus, this fungal genus might have an important role during plant naturalization to restrictive conditions. In relation to this, our previous studies of the microbiota that characterizes the soils from the Flooding Pampa in Argentina demonstrated that *Fusarium* is the dominant fungal genus in soils devoted to the promotion of *L. tenuis*, characterized by high salinity and alkalinity conditions (Nieva et al. 2018). In addition, the ability of the strains we isolated from roots of *Lotus* to tolerate different concentrations of NaCl might indicate the existence of a particular trait that let them to grow under unfavourable abiotic conditions in the soil. In support of this hypothesis, previous reports have demonstrated that many members of the genus *Fusarium* and other *Deuteromycetes* are tolerant to high NaCl concentrations (Tresner and Hayes 1971). Moreover, the ability of *F. solani* to tolerate severe alkaline conditions *in-vitro* has been previously demonstrated (Dumestre et al. 1997), and studies related to the distribution of fungi in alkaline soils have described several *Fusarium* species as alkali-tolerant and alkalophilic (Nagai et al. 1995). Thus, the combination of these abilities (tolerance to salinity and alkalinity) could have an important function in the ecosystem in favour of *Fusarium* species in natural environments, whereas the growth of other species is restricted because of the extreme environments.

Different studies have shown that fungal species classified as Dark Septate Endophytes (DSE), which have the particular attribute to growth and play important roles in extreme environments, confer interesting benefits to their hosts, such as phosphate acquisition and stress tolerance (Jumpponen and Trappe 1998). However, the absence of DSE from the microscopic observations of roots in our study (data non-shown) indicates that *Fusarium* species are the microorganisms that play this important role while they multiply both, inside the plant system and potentially in the soil.

### Plant growth promotion vs. Plant growth impairment

The inoculation of two *Lotus spp*. species with *F. solani* has shown differential effects on plant growth. Interestingly, despite of the phylogenetic proximity between *L. japonicus* and *L. tenuis* (Escaray et al. 2012), a contrasting effect was observed in the biomass of roots and shoots. Thus, growth promotion was evidenced in *L. japonicus* as a consequence of the infection. Similar effects in plant growth during symbiotic interactions with species from the genus *Fusarium* were observed previously in other plant species, such as *Zea mays* (Yates et al. 1997, Mehmood et al. 2019) and *Hordeum vulgare* (Maciá-Vicente et al. 2009). In turn, although *L. tenuis* did not show disease-related symptoms like chlorosis or wilt after fungal inoculation, an impairment of the growth was observed in this species. However, in spite of the negative effects on plant growth, we conceive that in this case *F. solani* could not be considered as a pathogen, because did not produce disease symptoms (Agrios 1988). It is possible that, under our experimental conditions, we are observing an intermediate stage of balance on the mutualism-pathogenesis *continuum* as proposed by Saikkonen (1998). The permanence of certain fungi inside the plant tissues, without the development of pathogenically symptoms, is related to the fungus capacity to remain inside the plant like a “latent pathogen”, till the change of the environmental conditions turn the balance to pathogenesis (Carroll 1988). In this trend, host specificity shown by some endophytes is an evolutionary trait (Kiers et al. 2010) and determines, in the long term, the transmission and persistence of the microorganism through plant generations.

The infection of *F. solani 142L52B* in the roots of *Lotus* plants was intercellular and limited to the cortex cells of roots. This kind of infections is in agreement with previous observations of other interaction between *Fusarium* endophytes and roots (Schulz et al. 2007). Considering this, it is plausible to think that this *Fusarium* endophyte may not manage to transmit infection vertically to plant offspring, so it can be classified in the *group IV* according to the classification proposed by Rodriguez (2009) for fungal endophytes. The endophytes belonging to this group are characterized by infections limited to the root tissues, and show a broad host range and horizontal transmission. It is of interest to note that there are not reports about *Fusarium* pathogens affecting *L. tenuis* in the Flooding Pampa to date. The absence of reports about pathogenic events in *L. tenuis* by these fungal species reinforces the idea that the mutualism between *Fusarium* and *Lotus* could play an important role in determining the biodiversity of the ecosystem and in plant fitness.

### Photosynthesis activity and energetic balance

According to previous studies, plant photosynthetic rates vary during fungal infections. For instance, it has been reported that mycorrhizal infections cause an increment in the photosynthetic rate in other legumes like *Glycine max* and *Vicia faba* (Kaschuk et al. 2009). By contrast, other reports indicated a decrease in plant photosynthesis as a consequence of the fungal infection by endophytes (Spiering et al. 2006). In the specific case of interactions established by species from *Fusarium*, previous works generally reported impairment in the PN, even when these infections were symptomless (Lorenzini et al. 1997, Pinto et al. 2000). In our experimental conditions, *L. japonicus* increased PN following *F. solani*infections. Importantly, as the PN value is measured through the rate of CO_2_ exchange; it results in an indirect measure of the rubisco carboxylation activity. In addition, the increment in the PN rate is in agreement with reduction in the levels of CO_2_ in the sub-stomatal cavity (C_i_), which indicate a higher rate of C fixation. Thus, these results indicate that CO_2_ getting into the sub-stomatal cavity is fixed more efficiently by rubisco in the photosynthetic process as a consequence of fungal infection.

The parameter Fv/Fm (Φ_Po_) was used previously to characterize the plant physiological status during pathogenic interactions (Barón et al. 2016, Pineda et al. 2018). Under our experimental conditions, the Fv/Fm values did not change among the treatments and *Lotus* species, indicating that the interactions could be considered non stressful for plants. Even more, the Fv/Fm values observed are in agreement with those obtained in non-stressed plants in previous studies conducted in two ecotypes of *L. japonicus*(Babuín et al. 2014, Campestre et al. 2016). Besides, the absence of statistically significant differences in other of the *OJIP*-derivated parameters between treatments in *L. japonicus* indicates that the structural components of the photosynthetic activity are not affected by the *F. solani* infection. The increase in the value of PN was associated to an increment in the concentration of sugars in *L. japonicus*. Similar effects were reported before as a result of the endophytic interactions between *Lolium perenne-Neotyphodium lolii* (Rasmussen et al. 2008) and *Lycopersicon esculentum-Leptodontidium orchidicola* (Andrade-Linares et al. 2011). In the case of *L. japonicus*, the amount of fructose and glucose in leaves correlated with the accumulation of its glycolytic intermediates fructose-6-phosphate and glucose-6-phosphate. In addition, the elevated levels in maltose can be attributed to an increased starch degradation rate in order to meet higher carbon requirements as a consequence of the interaction. In turn, the increase in the amount of glucose and fructose in roots demonstrates that carbohydrates may be readily transported to the root systems. These results reinforce the assumption that the fungal infection modified plant primary metabolism in order to supply more carbon to the invading fungus.

The differences observed in PIabs and additional *OJIP* parameters in *L. tenuis* indicate an alteration in the energetic status of the plants during infection. However, these modifications were not associated to a reduction in the value of PN. Interestingly, even though this species showed similar photosynthetic rates as *L. japonicus*, the amount of sugars detected in leaves were not modified by the infection, whereas the amount of sucrose decreases in roots. A significant reduction in the concentration of fructose, glucose and sucrose was also reported in the interaction between wheat and *F. moniliforme*, which was explained by the consumption of carbon resources by the endophyte located in roots (Bönnighausen et al. 2019). Whether a similar carbohydrate depletion mechanism explains the reduction of sugars and biomass in *L. tenuis* following fungal infection (Fig. 4) needs to be addressed.

On the other hand, it should be taken into account that sugars also function as signal molecules, which are involved in several plant processes such as growth and development (Rolland et al. 2006). Thus, hexoses as well as sucrose have been recognized as important signal molecules in source-sink regulation (Roitsch et al. 1999). Therefore, the differences in the concentration of hexoses in both plant species may be a consequence of the existence of complex and source-sink related signalling mechanisms that are differentially triggered in the two *Lotus* species during the infection process.

### Nutritional effects of fungal colonization

The fungal endophytes can provide nutritional benefits to the host, providing the plant with a better access to important elements such as N, P and K (Bolan et al. 1991). Nevertheless, our results indicate that there are no significant differences in the amount of N and P relative to dry biomass in roots and leaves following plant colonization in neither of the species evaluated. Thus, in spite of the phosphate solubilising ability of the *F. solani* strain used in this work, the increment of P and the slight decrease of N in leaves and roots of *L. japonicus* or *L. tenuis* were not significant (Fig. 8 a, b). These results indicate that phosphate facilitation could not be directly linked to the increment in the biomass in *L. japonicus*, and that plant growth enhancement of *L. japonicus* compared to *L. tenuis* could be due to a combination of several alternative mechanisms. In turn, we observed a significant increase of K in leaves of *L. tenuis* and a reduction in *L. japonicus*. This effect mirrors differential biomass accumulation in the two *Lotus* host species and suggests association of growth with a differential effect of *F. solani* on the transport and/or acquisition of K. The mechanisms by which *F. solani* infection, plant growth, and K accumulation are linked are currently unclear, but this macronutrient has been deeply studied in relation with the severity of different abiotic and biotic stresses in plants (Shabala and Pottosin 2014). Particularly, the effect of high K levels in crops systems has been related with reduced disease symptoms caused by *Fusarium spp*. (McClellan and Stuart 1947, Dastur et al. 1964, Schneider, 1985, Smiley et al. 1972). Moreover, K_2_SiO_3_ has been proposed as a useful agent to control pathogens (Chérif et al. 1994). Therefore, the possibility that the increase of K in *L. tenuis* could help to ameliorate the negative effects of *F. solani* may be a lead for future investigations of the *continuum* of beneficial to pathogenic plant microbe interactions.

### Concluding remarks

In this study we analysed the responses of two *Lotus* species to the infection by the strain *142L52B* of *F. solani* that was isolated from restrictive soils and showing a remarkable ability to solubilise phosphate and to tolerate salinity or alkaline conditions.

Our results demonstrate that *F. solani 142L52B is* able to colonize endophytically two *Lotus* species, *L. japonicus* and *L. tenuis*. The infections have differential consequences in the carbohydrate and photosynthesis status of both plant hosts. In the case of *L. japonicus*, we observed a growth promotion effect, which was associated to increased photosynthesis. In contrast, *L. tenuis* suffered growth impairment. In this case, even though the photosynthetic rate did not change, the functionality of PSII was affected and the amount of photoassimilates decreased. The reduction in the concentration of sugars could indicate differential carbohydrate utilization to support and differentially modulate or even limit by withholding nutrients (Schwachtje et al. 2018) the establishment of the plant-fungus interaction.

A better understanding of the mechanisms involved in the plant-fungal endophyte interactions will require complementary studies to detect the specific processes and signals that are involved in the switch between mutualism and pathogenesis.

## Acknowledgments

We thank Ines Fehrle for her technical support of the metabolomics analyses. This work was financial supported by the projects PICT 2014 and PICT 2015 funding by Agencia de Promoción Científica y Tecnológica, Argentina.

---

**Fig. S1** Number of leaves (a) and shoots (b) of *L. japonicus* and *L. tenuis* quantified at 37 days post inoculation with *F. solani 142L52B* (mean±standard deviation, n=10). Student’s test did not reveal significant differences at p<0.05.

**Supplementary material**. Sequences obtained by amplification of the regions ITS1-ITS2 and alpha-Transcription elongation factor of the 18S gene for the fungal strains selected according to the best performance to solubilise phosphate *in-vitro*.

## Abbreviations

ITS: : Internal Transcribed Spacer gene
TEF: : alpha Transcription Elongation Factor gene
PN: : Net photosynthesis
GS: : stomatal conductance
C_i_: : CO^2^ in the sub-stomatal cavity
PSII: : Photosystem II
*OJIP*: : O:Initial fluorescence, at 50 μs or less, J-I: intermediate levels at 2 ms and 30 ms, P: maximal fluorescence
Fv/Fm (Φ_Po,_): : Maximum quantum yield of primary photochemistry
PIabs: : Performance Index on the Absorption Basis
ABS: : Absorption flux
Sm: : normalized total complementary area above the *OJIP* transient
TRo: : Trapped energy flux
ETo: : Electron transport flux
DIo: : Dissipation flux
RC: : Reaction centre
CSo: : Cross section
Ψ_Eo_: : Electron transport from *quinone a* to plastoquinone efficiency
Φ_Eo_: : maximum yield of electron transport from *Qa* to plastoquinone primary photochemistry
GC-EI/TOF-MS: : gas chromatography coupled to electron impact ionization/time-of-flight mass spectrometry
N: : Total Nitrogen
P: : Total Phosphorus
K: : Total Potassium
DSE: : Dark septate endophytes.

## References

Agrios GN (2005) Introduction to plant pathology. Elsevier Academic Press Publication.

Andrade-Linares DR, Grosch R, Restrepo S, Krumbein A, Franken P (2011) Effects of dark septate endophytes on tomato plant performance. Mycorrhiza: 413–422.

Antonelli CJ, Calzadilla PI, Escaray FJ, Babuin MF, Campestre, MP, Rocco R, Bordenave CD, Perea García A, Nieva AS, Llames ME, Maguire V, Melani G, Sarena D, Bailleres M, Carrasco P, Paolocci F, Garriz A, Menendez A, Ruiz OA (2016) LOTUS spp: Biotechnological strategies to improve the Bioeconomy of lowlands in the Salado River Basin (Argentina). Agrofor, 1:43–53.

Antonelli CJ, Calzadilla PI, Vilas JM, Campestre MP, Escaray FJ, Ruiz OA (2019) Physiological and anatomical traits associated with tolerance to long-term partial submergence stress in the *Lotus* genus: responses of forage species, a model and an interspecific hybrid. Journal of Agronomy and Crop Science.

Babuin MF, Campestre MP, Rocco R, Bordenave CD, Escaray FJ, Antonelli C, Calzadilla P, Gárriz A, Serna E, Carrasco P, Ruiz OA, Menendez A (2014) Response to long-term NaHCO3-derived alkalinity in model Lotus japonicus Ecotypes Gifu B-129 and Miyakojima MG-20: transcriptomic profiling and physiological characterization. PloS one, 9(5), e97106.

Backhouse D, Burgess LW, Summerell BA (2001) Biogeography of *Fusarium*. In Fusarium. Paul E. Nelson Memorial symposium. APS Press, Bethesda, pp. 122–137.

Barón M, Pineda M, Pérez-Bueno ML (2016) Picturing pathogen infection in plants. Zeitschrift für Naturforschung C, 71: 355–368.

Berger S, Sinha AK, Roitsch T (2007) Plant physiology meets phytopathology: plant primary metabolism and plant–pathogen interactions. Journal of experimental botany 58: 4019–4026.

Berrocal-Lobo M, Molina A (2008) Arabidopsis defense response against Fusarium oxysporum. Trends in plant science 13: 145–150.

Bolan NS (1991) A critical review on the role of mycorrhizal fungi in the uptake of phosphorus by plants. Plant and soil 134: 189–207.

Bönnighausen J, Schauer N, Schäfer W, Bormann J (2019) Metabolic profiling of wheat rachis node infection by Fusarium graminearum–decoding deoxynivalenol-dependent susceptibility. New Phytologist: 459–469.

Bremner JM, Mulvaney CS (1982) Nitrogen—Total 1. Methods of soil analysis. Part 2. Chemical and microbiological properties, (methodsofsoilan2): 595–624.

Burke DW, Hall R (1991) Fusarium root rot. Compendium of bean diseases. Edited by R. Hall. American Phytopathological Society Press, St. Paul, Minn, 9–10.

Calzadilla, P. I., Maiale, S. J., Ruiz, O. A., & Escaray, F. J. (2016). Transcriptome response mediated by cold stress in Lotus japonicus. Frontiers in plant science, 7, 374.

Campestre MP, Antonelli C, Calzadilla PI, Maiale SJ, Rodríguez AA, Ruiz OA (2016) The alkaline tolerance in Lotus japonicus is associated with mechanisms of iron acquisition and modification of the architectural pattern of the root. Journal of plant physiology 206: 40–48.

Carroll G (1988) Fungal endophytes in stems and leaves: from latent pathogen to mutualistic symbiont. Ecology 69: 2–9.

Chérif M, Menzies JG, Ehret DL, Bogdanoff C, Belanger RR (1994) Yield of cucumber infected with *Pythium aphanidermatum* when grown with soluble silicon. HortScience 29: 896–897.

Dastur RH, Bhatt JG (1964) Relation of potassium to Fusarium wilt of flax. Nature 201, 1243.

Dethloff F, Erban A, Orf I, Alpers J, Fehrle I, Beine-Golovchuk O, Schmidt S, Schwachtje J, Kopka J (2014) Profiling methods to identify cold-regulated primary metabolites using gas chromatography coupled to mass spectrometry. Methods in Molecular Biology 1166: 171–197.

Di Pietro A, García-Maceira FI, Méglecz E, Roncero MIG (2001) A MAP kinase of the vascular wilt fungus *Fusarium oxysporum* is essential for root penetration and pathogenesis. Molecular microbiology 39: 1140–1152.

Douglas AE (2010) The symbiotic habit. Princeton University Press.

Dumestre A, Chone T, Portal J, Gerard M, Berthelin J (1997) Cyanide Degradation under Alkaline Conditions by a Strain of *Fusarium solani* Isolated from Contaminated Soils. Applied and environmental microbiology 63: 2729–2734.

Escaray FJ, Menendez AB, Gárriz A, Pieckenstain FL, Estrella MJ, Castagno LN, Ruiz OA (2012) Ecological and agronomic importance of the plant genus *Lotus*. Its application in grassland sustainability and the amelioration of constrained and contaminated soils. Plant Science, 182, 121–133.

Evans CGT, Herbert D, Tempest DW (1970) Chapter XIII the continuous cultivation of micro-organisms: 2. construction of a chemostat. In Methods in microbiology. Vol. 2, Academic Press, pp: 277–327.

García I, Mendoza R, Pomar MC (2008) Deficit and excess of soil water impact on plant growth of Lotus tenuis by affecting nutrient uptake and arbuscular mycorrhizal symbiosis. Plant and Soil, 304: 117–131.

Geiser DM, Jiménez-Gasco MM, Kang S, Makalowska I, Veeraraghavan N, Ward TJ, Zhang N, Kuldau GA, O’donnell K (2004). FUSARIUM-ID v. 1.0: a DNA sequence database for identifying *Fusarium*. European Journal of Plant Pathology, 110:473–479.

Handberg K, Stougaard J (1992) *Lotus japonicus,* an autogamous, diploid legume species for classical and molecular genetics. The Plant Journal 2: 487–496.

Hummel J, Strehmel N, Selbig J, Walther D, Kopka J (2010). Decision tree supported substructure prediction of metabolites from GC-MS profiles. Metabolomics 6: 322–333.

Imazaki I, Kadota I (2015) Molecular phylogeny and diversity of *Fusarium* endophytes isolated from tomato stems. FEMS microbiology ecology 91: fiv098.

Jumpponen A RI, Trappe JM (1998) Dark septate endophytes: a review of facultative biotrophic root-colonizing fungi. The New Phytologist 140: 295–310.

Kaschuk G, Kuyper TW, Leffelaar PA, Hungria M, Giller KE (2009) Are the rates of photosynthesis stimulated by the carbon sink strength of rhizobial and arbuscular mycorrhizal symbioses?. Soil Biology and Biochemistry 41:1233–1244.

Kavroulakis N, Ntougias S, Zervakis GI, Ehaliotis C, Haralampidis K, Papadopoulou KK (2007) Role of ethylene in the protection of tomato plants against soil-borne fungal pathogens conferred by an endophytic *Fusarium solani* strain. Journal of Experimental Botany 58: 3853–3864.

Kirkbride JH (2006) The scientific name of narrow-leaf trefoil. Crop science, 46: 2169–2170.

Kuldau GA, Yates IE (2000) Evidence for *Fusarium* Endophytes. Microbial endophytes: 85.

Kusari S, Zühlke S, Spiteller M (2009) An endophytic fungus from *Camptotheca acuminata* that produces camptothecin and analogues. Journal of Natural Products 72: 2–7.

Luedemann A, Strassburg K, Erban A, Kopka J (2008) TagFinder for the quantitative analysis of gas chromatography—mass spectrometry (GC-MS)-based metabolite profiling experiments. Bioinformatics 24: 732–737.

Lisec J, Schauer N, Kopka J, Willmitzer L, Fernie AR (2006) Gas chromatography mass spectrometry–based metabolite profiling in plants. Nature protocols 1:387.

Lorenzini G, Guidi L, Nali C, Ciompi S, Soldatini GF (1997) Photosynthetic response of tomato plants to vascular wilt diseases. Plant science 124: 143–152.

Lyons PC, Evans JJ, Bacon C W (1990) Effects of the fungal endophyte Acremonium coenophialum on nitrogen accumulation and metabolism in tall fescue. Plant physiology 92: 726–732.

Maciá-Vicente JG, Jansson HB, Abdullah SK, Descals E, Salinas J, Lopez-Llorca LV (2008) Fungal root endophytes from natural vegetation in Mediterranean environments with special reference to *Fusarium spp*. FEMS microbiology ecology 64: 90–105.

Maciá-Vicente JG, Rosso LC, Ciancio A, Jansson HB, Lopez-Llorca LV (2009) Colonisation of barley roots by endophytic *Fusarium equiseti* and *Pochonia chlamydosporia:* effects on plant growth and disease. Annals of Applied Biology 155: 391–401.

Matuo T, Snyder WC (1973) Use of morphology and mating populations in the identification of formae speciales in *Fusarium solani*. Phytopathology 63: 562–565.

McClellan WD, Stuart NW (1947) The influence of nutrition on *Fusarium* basal rot of narcissus and on *Fusarium* yellows of gladiolus. American journal of botany 34: 88–93.

McMullen M, Jones R, Gallenberg D (1997) Scab of wheat and barley: a re-emerging disease of devastating impact. Plant disease, 81: 1340–1348.

Mehmood A, Hussain A, Irshad M, Hamayun M, Iqbal A, Rahman H, Tawab A, Ahmad A, Ayaz S (2019) Cinnamic acid as an inhibitor of growth, flavonoids exudation and endophytic fungus colonization in maize root. Plant Physiology and Biochemistry 135: 61–68.

Murphy J. Riley J P (1962)A modified single solution method for the determination of phosphate in natural waters. Analytica chimica acta 27: 31–36.

Nagai K, Sakai T, Rantiatmodjo RM, Suzuki K, Gams W, Okada G (1995) Studies on the distribution of aikalophilic and alkali-tolerant soil fungi I. Mycoscience 36: 247–256.

Nieva AS, Bailleres MA, Llames ME, Taboada MA, Ruiz OA, Menéndez A (2018) Promotion of Lotus tenuis in the Flooding Pampa (Argentina) increases the soil fungal diversity. Fungal Ecology 33: 80–91.

Nilsson RH, Larsson KH, Taylor AFS, Bengtsson-Palme J, Jeppesen TS, Schigel D, Kennedy P, Picard K, Glöckner FO, Tedersoo L, Saar I, Kõljalg U, Abarenkov K (2018) The UNITE database for molecular identification of fungi: handling dark taxa and parallel taxonomic classifications. Nucleic Acids Research, DOI: 10.1093/nar/gky1022.

Phillips JM, Hayman DS (1970) Improved procedures for clearing roots and staining parasitic and vesicular-arbuscular mycorrhizal fungi for rapid assessment of infection. Transactions of the British mycological Society 55: 158–161.

Pineda M, Pérez-Bueno ML, Barón M (2018) Detection of bacterial infection in melon plants by classification methods based on imaging data. Frontiers in plant science 9: 164.

Pinto LSRC, Azevedo JL, Pereira JO, Vieira MLC, Labate CA (2000) Symptomless infection of banana and maize by endophytic fungi impairs photosynthetic efficiency. The New Phytologist 147: 609–615.

Rasmussen S, Parsons AJ, Fraser K, Xue H, Newman JA (2008) Metabolic profiles of Lolium perenne are differentially affected by nitrogen supply, carbohydrate content, and fungal endophyte infection. Plant physiology 146: 1440–1453.

Richardson MD, Hoveland CS, Bacon CW (1993) Photosynthesis and stomatal conductance of symbiotic and nonsymbiotic tall fescue. Crop science 33: 145–149.

Rodriguez RJ, White JrJF, Arnold AE, Redman ARA (2009) Fungal endophytes: diversity and functional roles. New phytologist 182: 314–330.

Roitsch T (1999) Source-sink regulation by sugar and stress. Current opinion in plant biology: 198–206.

Rolland F, Baena-Gonzalez E, Sheen J (2006) Sugar sensing and signaling in plants: conserved and novel mechanisms. Annu. Rev. Plant Biol: 675–709.

Romberg MK, Davis RM (2007) Host range and phylogeny of *Fusarium solani f. sp. eumartii* from potato and tomato in California. Plant Disease 91: 585–592.

Roncero MIG, Hera C, Ruiz-Rubio M, Maceira FIG, Madrid MP, Caracuel Z, Calero F, Delgado-Jarana J, Roldán-Rodriguez R,Martinez AL,Velasco C, Roa J, Martinez-Urdiroz M, Córdoba D, Di Pietro A (2003) Fusarium as a model for studying virulence in soilborne plant pathogens. Physiological and Molecular Plant Pathology 62: 87–98.

Ruan Y, Kotraiah V, Straney DC (1995) Flavonoids stimulate spore germination in *Fusarium solani* pathogenic on legumes in a manner sensitive to inhibitors of cAMP-dependent protein kinase. MPMI- Molecular Plant Microbe Interactions 8: 929–938.

Saikkonen K, Faeth SH, Helander M, Sullivan TJ (1998) Fungal endophytes: a continuum of interactions with host plants. Annual review of Ecology and Systematics 29: 319–343.

Saikkonen K, Gundel PE, Helander M (2013) Chemical ecology mediated by fungal endophytes in grasses. Journal of chemical ecology 39: 962–968.

Schneider RW (1985) Suppresiion of Fusarium yellows of celery with potassium, chloride, and nitrate. Phytopathology 75: 40–48.

Schulz BJ, Boyle CJ, Sieber TN (2007) Microbial root endophytes (Vol. 9). Springer Science & Business Media.

Schwachtje J, Fischer A, Erban A, Kopka J (2018) Primed primary metabolism in systemic leaves: a functional systems analysis. Scientific Reports 8: 216.

Shabala S, Pottosin I (2014) Regulation of potassium transport in plants under hostile conditions: implications for abiotic and biotic stress tolerance. Physiologia Plantarum 151: 257–279.

Shweta S, Zuehlke S, Ramesha BT, Priti V, Kumar PM, Ravikanth G, Spiteller R, Vasudeva R, Shaanker, RU (2010) Endophytic fungal strains of *Fusarium solani,* from *Apodytes dimidiata* E. Mey. ex Arn (Icacinaceae) produce camptothecin, 10-hydroxycamptothecin and 9-methoxycamptothecin. Phytochemistry 71: 117–122.

Smiley RW, Cook RJ, Papendick RI (1972) *Fusarium* foot rot of wheat and peas as influenced by soil applications of anhydrous ammonia and ammonia potassium azide solutions. Phytopathology 62: 86.

Spiering MJ, Greer DH, Schmid JAN (2006) Effects of the fungal endophyte, Neotyphodium lolii, on net photosynthesis and growth rates of perennial ryegrass *(Lolium perenne)* are independent of in planta endophyte concentration. Annals of botany 98: 379–387.

Tayung K, Barik BP, Jha DK, Deka DC (2011) Identification and characterization of antimicrobial metabolite from an endophytic fungus, *Fusarium solani* isolated from bark of Himalayan yew. Mycosphere 2: 203–213.

Toby Kiers E, Palmer TM, Ives AR, Bruno JF, Bronstein JL (2010) Mutualisms in a changing world: an evolutionary perspective. Ecology letters 13: 1459–1474.

Tresner HD, Hayes JA (1971) Sodium chloride tolerance of terrestrial fungi. Applied Microbiology, 22: 210–213.

Vilas JM, Romero FM, Rossi FR, Marina M, Maiale SJ, Calzadilla PI, Pieckenstain FL, Ruiz OA, Gárriz A (2018) Modulation of plant and bacterial polyamine metabolism during the compatible interaction between tomato and Pseudomonas syringae. Journal of plant physiology 231: 281–290.

White TJ, Bruns T, Lee SJWT, Taylor JL (1990) Amplification and direct sequencing of fungal ribosomal RNA genes for phylogenetics. PCR protocols: a guide to methods and applications 18: 315–322.

Xu M, Gessner G, Groth I, Lange C, Christner A, Bruhn T, Deng Z, Li X, Heinemann H, Grabley S, Bringmann G, Sattler I, Lin W (2007) Shearinines D–K, new indole triterpenoids from an endophytic Penicillium sp.(strain HKI0459) with blocking activity on large-conductance calcium-activated potassium channels. Tetrahedron 63: 435–444.

Yates IE, Bacon CW, Hinton DM (1997) Effects of endophytic infection by *Fusarium moniliforme* on corn growth and cellular morphology. Plant disease 81: 723–728.

