## Supplementary figures and images for "The fungal endophyte *Fusarium solani* provokes differential effects on the energy balance of two *Lotus* species"

### Supplemental Figure 1

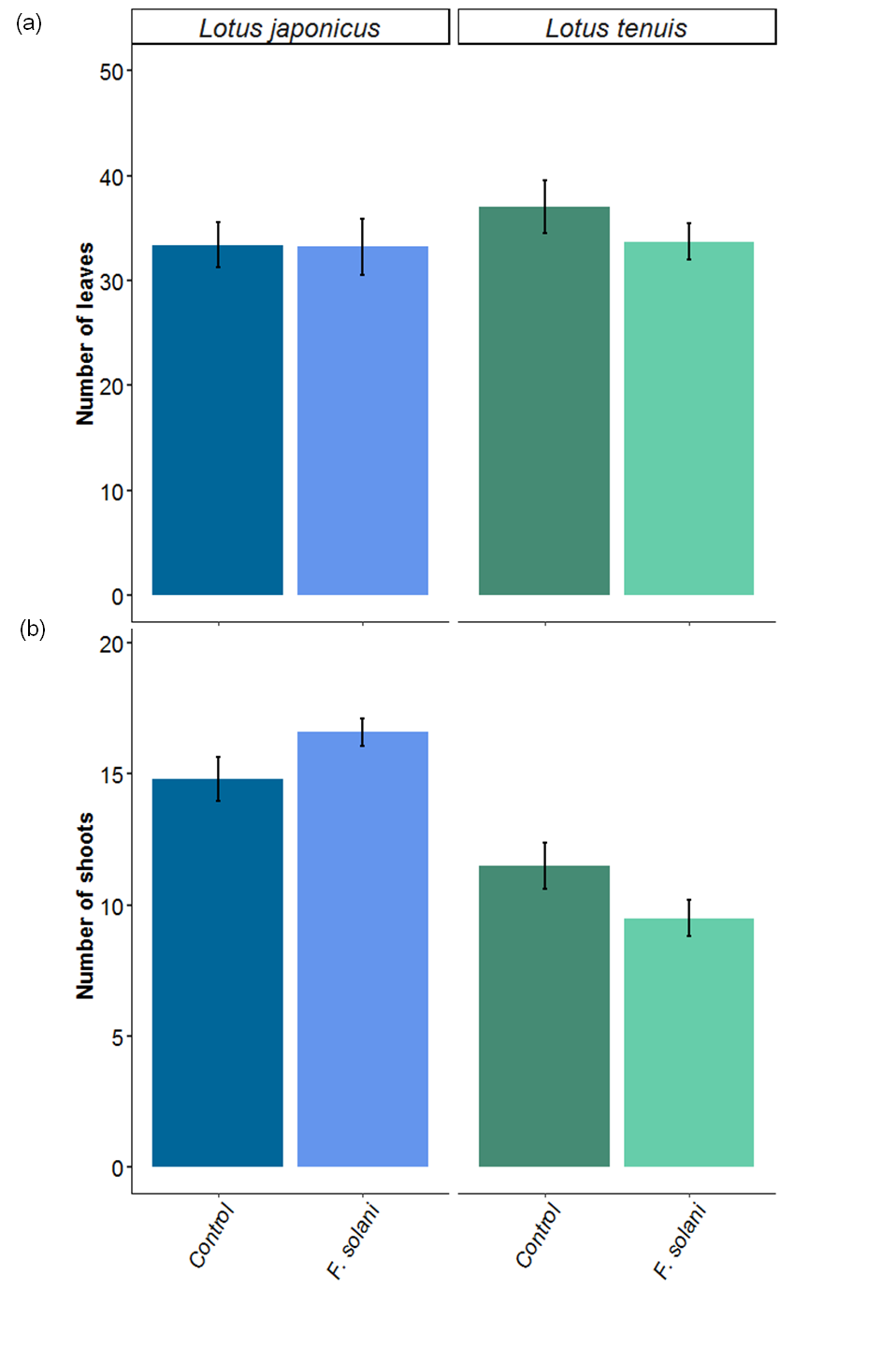
