## Supplementary material for "The fungal endophyte *Fusarium solani* provokes differential effects on the energy balance of two *Lotus* species"

**Supplementary material .** Sequences obtained by amplification of the regions ITS1-ITS2 and alpha-Transcription elongation factor of the 18S gene for the fungal strains selected according to the best performance to solubilise phosphate *in-vitro*.

>Fusarium_oxysporum_142L52A_ITS1-ITS2

AACAAGGTCTCCGTTGGTGAACCAGCGGAGGGATCATTACCGAGTTTACAACTCCCAAACCCCTGTGAACATACCACTTGTTGCCTCGGCGGATCAGCCCGCTCCCGGTAAAACGGGACGGCCCGCCAGAGGACCCCTAAACTCTGTTTCTAATATGTAACTTCTGAGTAAAACCATAAATAAATCAAAACTTTCAACAACGGATCTCTTGGTTCTGGCATCGATGAAGAACGCAGCAAAATGCGATAAGTAATGTGAATTGCAGAATTCAGTGAATCATCGAATCTTTGAACGCACATTGCGCCCGCCAGTATTCTGGCGGGCATGCCTGTTCGAGCGTCATTTCAACCCTCAAGCACAGCTTGGTGTTGGGACTCGCGTTAATTCGCGTTCCTCAAATTGATTGGCGGTCACGTCGAGCTTCCATAGCGTAGTAGTAAAACCCTCGTTACTGGTAATCGTCGCGGCCACGCCGTTAAACCCCAACTTCTGAATGTTGACCTCGGATCAGGTAGGAATACCCGCTGAACTTAAGCAT

>Fusarium_solani _142L52B_ITS1-ITS2

TACCGAGTTATACAACTCATCAACCCTGTGAACATACCTATAACGTTGCCTCGGCGGGAACAGACGGCCCCGTAACACGGGCCGCCCCCGCCAGAGGACCCCCTAACTCTGTTTCTATAATGTTTCTTCTGAGTAAACAAGCAAATAAATTAAAACTTTCAACAACGGATCTCTTGGCTCTGGCATCGATGAAGAACGCAGCGAAATGCGATAAGTAATGTGAATTGCAGAATTCAGTGAATCATCGAATCTTTGAACGCACATTGCGCCCGCCAGTATTCTGGCGGGCATGCCTGTTCGAGCGTCATTACAACCCTCAGGCCCCCGGGCCTGGCGTTGGGGATCGGCGGAAGCCCCCTGCGGGCACAACGCCGTCCCCCAAATACAGTGGCGGTCCCGCCGCAGCTTCCATTGCGTAGTAGCTAACACCTCGCAACTGGAGAGCGGCGCGGCCACGCCGTAAAACACCCAACTTCTGAATGTTGACCTCGAATCAGGTAGGAATACCCGCTGAACTTAAGCATATCAAA

>Fusarium_solani_142L52B_alpha_TEF1

CCCCGCCTGGCATCTCGGGCGGGGTATTCATCAGTCACTTCATGCTGACAATCATCGACAGACCGGTCACTTGATCTACCAGTGCGGTGGTATCGACAAGCGAACCATCGAGAAGTTCGAGAAGGTTGGTGACATCTCCCCCGATCGCGCCTTGCTATTCCACATCGAATTCCCTCCCTCGCGATACGCTCTGCGCCCGCTTCTCCCGAGTCCCAAAATTTTTGCGGTCCGACCGTAATTTTTTTTTGGTGGGGCTTTTACCCCGCCACTCGGGCGAGTAAGGAGGACAAGACCAGCGGCGAGGATGCAAGCGGGAAGAAACCCTCTTGGCGCGCATCATCACGTGGTTCACAACAGACGCTAACCGGTCCAACAATAGGAAGCCGCTGAGCTCGGTAAGGGT

>Fusarium_oxysporum_142L52A_alpha_TEF1

TGGGTAAGGAGGACAAGACTCACCTTAACGTCGTCGTCATCGGCCACGTCGACTCTGGCAAGTCGACCACTGTGAGTACTCTCCTCGACAATGAGCATATCTGCCATCGTCAATCCCGACCAAGACCTGGCGGGGTATTTCTCAAAGTCAACATACTGACATCGTTTCACAGACCGGTCACTTGATCTACCAGTGCGGTGGTATCGACAAGCGAACCATCGAGAAGTTCGAGAAGGTTAGTCACTTTCCCTTCAATCGCGCGTCCTTTGCCCATCGATTTCCCCTACGACTCGAAACGTGCCCGCTACCCCGCTCGAGACCAAAAATTTTGCAATATGACCGTAATTTTTTTGGTGGGGCACTTACCCCGCCACTTGAGCGAAGGGAGCGTTTGCCCTCTTACCATTCTCACAACCTCAATGAGTGCGTCGTCACGTGTGAAGCAGT
